# Intragenomic homogeneity on Liberibacter 16S rDNA confirms phylogeny and explains ecological strategy

**DOI:** 10.1101/016188

**Authors:** Warrick R. Nelson, Nelson A. Wulff, Joseph M. Bové

## Abstract

Three of the five currently recognized “*Candidatus* Liberibacter” spp., “*Ca*. L. asiaticus” (Las), “*Ca*. L. americanus” (Lam) and “*Ca*. L. solanacearum”, and the newly erected genus *Liberibacter* species, L. *crescens* (Lcr), have had their genomes sequenced. In all four cases there are three homogeneous copies of the 16S rRNA gene, one present as the reverse complement of the other two. 16S intragenic homogeneity is common within the α-Proteobacteria. The presence of three 16S rRNA copies indicates an advantage for a rapid response of population increase to favourable growth conditions. The metabolic cost of carrying multiple copies is avoided during periods of low cellular activity as this situation occurs at low temperatures, for example overwintering in deciduous plants or in a dormant insect host.

A large insertion in the 16S rDNA sequence of three species compared to the other three species indicates a dichotomy in the *Liberibacter* genus and provides a phylogenetic signal of closeness to the proximal node within the Rhizobiaceae. In spite of similar symptoms in *Citrus* crops associated with Lam and Las infections, these species belong on either side of this dichotomy, thus confirming Lam as phylogenetically closer to the proximal node than Las.

## Introduction

The bacterial 16S rRNA gene is highly conserved and thus serves as a very useful marker of evolutionary relatedness in phylogenetic studies [1]. Bacterial species may contain one or more copies of this gene. A single copy is common where bacteria need to conserve resources as there is a metabolic cost associated with multiple copies, for example in an environment poor in nutrients and thus limiting growth potential [2]. Multiple homogeneous copies occur in species adapted to environments where speed of multiplication is important because of only temporarily good growth conditions [3]. Concerted evolution generally maintains these as homogeneous copies, probably because of a metabolic cost associated with combined expression of heterogeneous copies [4]. Multiple but heterogeneous copies have been recorded where the 16S gene copies are so different as to be able to function under very different conditions, especially particularly high or low temperatures, thus adapting the bacteria to grow across a broader temperature range than normal [5]. However, because the 16S gene is so commonly used as a phylogenetic marker, intragenic heterogeneity has resulted in an overestimation of bacterial species diversity [6, 7].

Liberibacter species are new to science, commonly in association with significant crop damage, especially of citrus, potato, tomato, carrot, celery and their close relatives [8]. They are members of the Rhizobiaceae family in the α division of Proteobacteria [9] and are described as obligate intracellular endogenous bacteria with an alternating insect (psyllid)/plant host life cycle. The insect/plant host species composition varying primarily by liberibacter species [8]. The earliest speciation event by molecular clock technique is 309 Ma [10], supported by biogeography to a putative Pangaean origin [11]. Six species have so far been discovered and named.

The genus “*Candidatus* Liberibacter” and the two species “*Ca*. L. africanus” (Laf) and “*Ca*. L. asiaticus” (Las), associated with huanglongbing (HLB) disease in citrus crops, were first described by Jagoueix, Bové and Garnier [9]. While the genome of Laf has not yet been sequenced, the Las genome sequence has been obtained from five different sources [12–16]. Comparison of a large number of Las 16S rDNA sequences indicates that there is a single haplotype in this species [17, 18] in spite of the very wide geographic spread it has now reached [19]. Las has been reported to exist in a number of genetic groupings, depending on the DNA segment analysed [20] and the divergence on other DNA segments may be more useful for epidemiological purposes. Las is also known from close citrus relatives commonly grown for ornamental and culinary purposes, especially *Murraya* species [21–23].

“*Ca*. L. americanus” (Lam) is also associated with HLB in citrus crops [24], as well as infecting *Murraya paniculata* [25]. The Lam genome has been sequenced [26]. At present this species has only been reported from Brazil; single reports from China and Texas (USA) have not been confirmed.

“*Ca*. L. solanacearum” (Lso) [27] (syn “*Ca*. L. psyllaurous”) [28] is associated with diseases of Solanaceae crops (especially potato and tomato) in North America and Apiaceae crops (especially carrot and celery) in Europe [29] and North Africa [30]. Five haplotypes with geographic and psyllid/plant host differences have been described, designated LsoA, B, C, D and E [31–33]. The genome has been sequenced [34], most likely the B haplotype.

*Liberibacter crescens* (Lcr), as the only cultured species within the liberibacters, is the first representative of the new genus *Liberibacter* to be validly published and named [35]. This species has been isolated from expressed sap of a defoliating mountain papaya in Puerto Rico [35]. The insect phase is currently unknown. The *L. crescens* genome has been sequenced [36].

Taxonomic errors may occur in the presence of intragenomic heterogeneity on the 16S rRNA gene [7] and it is known that liberibacter genomes may hold more than one 16S gene copy [37]. By combining knowledge of the 16S gene copy number and heterogeneity, phylogenetic status and some clear pointers to the evolutionary responses to the intracellular conditions can be deduced for the liberibacter species.

## Materials and Methods

*Liberibacter* genomes were identified in the GenBank database (http://www.ncbi.nlm.nih.gov/genome/?term=liberibacter), and the 16S rDNA sequences downloaded after identifying the genes through the sequence annotations. These annotations also indicate position of the gene within the genome, orientation and length, summarised in Table 1. Sequences were aligned in ClustalX v2.1 [38].

**Table 1.**
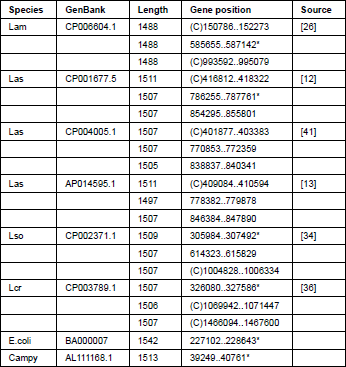
16S rDNA operons and nucleotide numbers within the genome. * indicates operon used for thermal stability curves calculated with RNAdraw

| Species | GenBank | Length | Gene position | Source |
| --- | --- | --- | --- | --- |
| Lam | CP006604.1 | 1488 | (C)150786..152273 | [26] |
|  |  | 1488 | 585655..587142* |  |
|  |  | 1488 | (C)993592..995079 |  |
| Las | CP001677.5 | 1511 | (C)416812..418322 | [12] |
|  |  | 1507 | 786255..787761* |  |
|  |  | 1507 | 854295..855801 |  |
| Las | CP004005.1 | 1507 | (C)401877..403383 | [41] |
|  |  | 1507 | 770853..772359 |  |
|  |  | 1505 | 838837..840341 |  |
| Las | AP014595.1 | 1511 | (C)409084..410594 | [13] |
|  |  | 1497 | 778382..779878 |  |
|  |  | 1507 | 846384..847890 |  |
| Lso | CP002371.1 | 1509 | 305984..307492* | [34] |
|  |  | 1507 | 614323..615829 |  |
|  |  | 1507 | (C)1004828..1006334 |  |
| Lcr | CP003789.1 | 1507 | 326080..327586* | [36] |
|  |  | 1506 | (C)1069942..1071447 |  |
|  |  | 1507 | (C)1466094..1467600 |  |
| E.coli | BA000007 | 1542 | 227102..228643* |  |
| Campy | AL111168.1 | 1513 | 39249..40761* |  |

Minimal free energy and the thermal stability curves were calculated using RNAdraw v1.1b2 [39] with the specific sequences used indicated in Table 1. *Escherichia coli* (E.coli GenBank BA000007) and *Campylobacter jejuni* (Campy GenBank AL111168.1) were added for comparison as they are also Proteobacteria but in the γ and ε divisions respectively.

## Results

The genomes of all four species of liberibacter, Las, Lso, Lam and Lcr, contain three copies of the 16S rRNA gene with one copy as the reverse complement of the other two. Aligning all three copies per genome retained the clear species separation as previously reported [11], with almost complete intragenomic homogeneity. The main differences were at the ends, most likely representing difficulty in determining the up- and down-stream promoter regions. A few single nucleotide polymorphisms (SNPs), indicating minor heterogeneity between copies, are more likely to be sequencing errors as a comparison at species level against many other accessions from GenBank do not repeat these SNPs (data not shown).

Although several rearrangements have taken place in liberibacter genomes [26, 36, 42], a high degree of gene conservancy is observed in the vicinity of ribosomal operons in all liberibacter genomes sequenced to date. If we order all genomes starting from the *dna*A gene, this conservancy is clearly shown in Figure 1. For instance, the three ribosomal operons of Lcr have the same orientation as those of the phylogenetically close Lam [35]; similarly, for Lso and Las, the orientation is also the same [11], but different from the Lcr/Lam orientation. In addition, mrp is located immediately downstream of the 5SrRNA gene 10 times out of 12, irrespectively of ribosomal operon orientation (compare operon 5 and 9, 6 and 8, as well as 7 and 12, 9 and 11).

**Figure 1.**
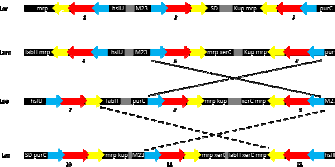
Orientation of rrn taken from Lcr, Lam, Lso and Las genomes, assuming rrn starting from dnaA gene, as observed in Lam [26]. Dotted lines between Lam/Lcr and Lso indicates rrn inversion, with second and third rrn operons being inverted in Lso. Dashed lines between Lso and Las indicates rrn inversion, with the first rrn being inverted and located as the third rrn operon in Las. Arrows indicatesBlue arrow = 16SrDNA; Red arrow = 23SrDNA and yellow arrow = 5SrDNA. purC (phosphoribosylaminoimidazole succinocarboxamide synthase), mrp (multiple resistance and pH adaptation), xerC gene (site-specific tyrosine recombinase), fabH gene (3-oxoacyl-[acyl III), kup (potassium uptake transporter), hslU (ATP-binding subunit from the ATP dependent protease); M23 (peptidase M23), SD (superoxide dismutase). Numbers in italics indicate rrn operon. Not to scale.

Thermal stability curves show similarities across the liberibacter species with marked differences to E.coli and Campy (Figure 2). Even more marked are the differences at higher temperat ure ranges for DNA derived from thermotolerant extremophiles. However, possibly more interesting is the almost complete lack of differences in the 10–40°C range (Figure 2).

**Figure 2.**
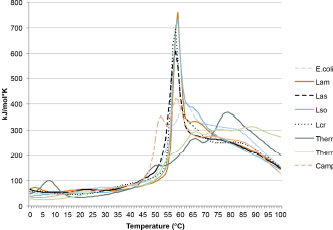
Predicted thermal stability curves of 16S rDNA from four liberibacter spe plus E.coli and Campy for comparison, the specific copy illustrated is indicated in Table 1, as well as *Thermoanaerobacter tengkongensis* (GenBank AE008691) 16S operons ThermoA and ThermoC [5] illustrating possible differences of rRNA bacterial species.

Comparing the 16S rRNA genes of the Lso genome sequenced (Table 1, Lso CP002371.1) with other Lso 16S accessions, indicates that the sequenced genome corresponds to haplotype LsoB, although it is not specifically identified as such. This haplotype has been associated withsymptomatic solanaceous crops of Central USA, East of the Rocky Mountains [31]. One of the copies from the sequenced genome has two SNPs, indicating LsoA haplotype. This could suggest that the source of liberibacter for genome assembly was a mixed haplotype infection, reinforced by the SNPs noted earlier in this species on the 23S and 5S genes. Mixed infections with these two haplotypes have been reported [43].

Although the Lcr gene is reported as longer than the Lam version, closer inspection of the sequences indicates no other insertions, but rather variation in reporting at both the 5’ and 3’ ends accounts for the discrepancy.

A large insertion of up to 14 nt occurs in Las and Lso (Figure 3). This insertion is positioned at nucleotides 1141/1142 of E.coli 16S rRNA gene. In E.coli this nucleotide locates inside helix 39 [44], which interacts with proteins S10 and S9, also present in liberibacter genomes. That this is an insertion event in Las and Lso is illustrated by the lack of these nucleotides in related Rhizobiaceae species as well as the more distantly related Proteobacteria species.

**Figure 3.**
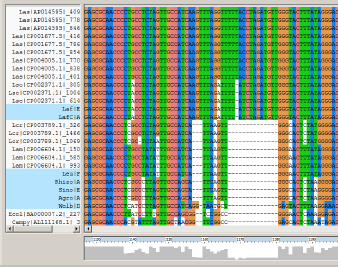
Partial screen capture of the ClustalX alignment showing the 14/13bp length insertion in the Las/Lso sequences compared to Lam/Lcr and E.coli/Campy. Other α-species are highlighted including two other liberibacter species, “*Ca*. L. africanus” (Laf, GenBank EU921619) and subspecies capensis (LafC, GenBank. L europaeus” (Leu, GenBank FN678792), the more distantly related *bium leguminosarum* (Rhizo, GenBank AY509900), (Sino, GenBank EU286550), *Agrobacterium tumefaciens* proteobacterium from the Ricketsiaceae “*Ca*. Wolbachia inokumae” (Wolb, GenBank DQ402518).

## Discussion

Intragenomic homogeneity of multiple 16S operons within α-proteobacteria is common [45] and liberibacter species fit this pattern. Concerted evolution is a driver towards homogeneity as heterogeneity comes at a metabolic cost [4]. The metabolic cost of heterogeneity is of lesser evolutionary importance only where other drivers occur, such as adaptation to extremes of temperature as illustrated in some extremophiles, for instance *Thermoanaerobacter* species [5].

Most bacteria possess one or two copies of the 16S gene, with three or more copies being considerably less common [7]. Proteobacteria generally have an average copy number of 3.94, but removing the much higher number common in the γ division results in fewer than 3 copies for the rest of the proteobacteria [45]. Thus the three copies in the liberibacters is towards the high side for a species in the α division.

Some studies suggest that liberibacters face competition within their plant phloem environment [46, 47] and their insect host microbial biome [48]. Retention of a relatively high number of 16S copies suggests an evolutionary advantage of a rapid increase in population in response to an improved environment for reproduction. However, a more detailed microbial diversity analysis specifically of phloem cells suggests a very limited microbial competition for liberibacter [49]. The many studies using electron microscopy easily identify liberibacter bacteria in symptomatic tissues of infected plants and no or fewer liberibacters in the phloem of infected, but yet asymptomatic plants; bacteria other than liberibacters are not detected in the phloem of infected plants. The evolutionary advantage for liberibacter species of multiple 16S copies therefore does not derive from competition with other bacterial species. Possibly the advantage derives from relatively short periods of high nutrient availability during the main growing season of both plant and insect phases. This would suggest strong seasonal differences in growth rates of these hosts.

However, the metabolic cost of the copy number may play a role as evidenced by apparently low titre in plant hosts during periods of low growth but higher titre during growth flushes of citrus [50, 51] or after storage in seed tubers of potatoes [52, 53]. An extreme of survival in root systems, even after the tree has been killed [54], indicates the potential for long term survival even in a dormant host. Deciduous plants exhibit winter dormancy and presumably the metabolic cost to liberibacter of multiple rRNA genes is less important at low temperatures common in these conditions. This suggests an evolutionary advantage of a higher copy number to increase population size rapidly when growth conditions are favourable while the metabolic costs are reduced because of low growth conditions for liberibacter in the dormant plant and/or insect phases.

Liberibacter species are known to be relatively intolerant of high temperatures, with Las being regarded as the most tolerant. Las can be eliminated from plants exposed to 42°C [55] while Lam and Lso were not transmitted to host plants at temperatures above 32°C [56, 57]. Thermal stability curves for 16S (Figure 2), while suggesting minor differences between species, do not indicate the major differences in thermal stability of intragenomically divergent 16S copies in thermophilic extremophiles such as in *Thermoanaerobacter* [5]. For example, the thermal stability curves for Campy and E.coli also do not indicate that the 16S gene is involved in determining growth temperatures, Campy has a growth range of 30–46°C [58] and E.coli an optimum of 37°C [59]. There is a danger of anticipating that intergenomic heterogeneity of 16S copies is associated with sensitivity to different temperatures. These observations suggest that this might only be the case for some extreme thermophilic bacteria and differences in optimal and survival temperature responses in liberibacter species are not as a result of 16S gene sensitivity.

Heat sensitivity suggests a cooler climate requirement, yet some of the regions where liberibacters are a problem in crops are not cool during the summer months. If foliage in these hotter periods becomes too hot for liberibacter survival, presumably the bacteria survive within the plants in cooler plant parts, especially in below ground parts, such as roots and tubers. This could explain the observation that plant sampling for liberibacter detection appears to be more reliable from roots or low down on the plant [51, 60].

Assuming that the small number of SNPs between copies in the same genome represent sequencing ambiguities or errors rather than true intragenomic heterogeneity, there has clearly been sufficient time for concerted evolution to be effective after species and haplotype separation. Although it is commonly stated that genomic variation is accelerated in obligate endosymbionts, this is unlikely to be true of what is effectively a clonal system on the highly conserved 16S gene and within the highly ecologically constrained biome of these obligate intracellular endogenous bacteria, as opposed to genomic loss of genes coding for proteins available from the host environment. Thus we can have simultaneously large and rapid changes at the genome level combined with almost no variation on highly conserved genes such as 16S, hence the value of the 16S gene for taxonomic purposes [1]. No changes in 16S rRNA genes and also no major changes in upstream associated ORFs (Figure 1) support genomic variation largely by gene loss rather than gene rearrangements.

The large insertion event representing about 0.9% of the gene (Figure 3) occurs within Las and Lso genomes but not in Lam or Lcr. Adding 16S sequences for the other liberibacter species but without genome sequences available (Laf, Leu) as well as a number of other Rhizobiaceae and more distantly related proteobacterial species, demonstrates this is an insertion event in the precursor species prior to Las and Lso speciation. This observation confirms that Lam is more closely positioned phylogenetically to the proximal node of the Rhizobiaceae than Las or Lso [40]. Clearly, the species form two phylogenetic clusters, those evolving prior to this insertion (Lam, Lcr, Leu) and those evolving from a common ancestor after this large insertion event (Laf, Las, Lso). This phylogenetic clustering is also supported by the gene arrangements and inversions illustrated in Figure 1, thus a dichotomy occurs within liberibacter species.

## Conclusion

The liberibacter species, whose genomes have been sequenced, universally contain three copies of essentially homogeneous 16S rRNA genes, and thus 16S-based phylogenies are sound. This copy number implies an evolutionary advantage of speed of population response over the metabolic cost associated with more copies. This suggests an ecosystem requiring speed to optimise growth in short periods rather than competing against other bacterial species, most likely to occur when conditions suitable for population growth are erratic. The lower evolutionary importance associated with metabolic cost of more rRNA gene copies implies that periods of low nutrient availability coincide with periods when cell division does not occur (low metabolic activity in bacteria). This suggests that the dormant phase of either insect or plant hosts is associated with low temperatures and therefore the disadvantage of high copy number is avoided.

